# Relationship between concentration of cytokinin for efficient shoot regeneration and seed coat color in leaf lettuce

**DOI:** 10.1101/2024.12.10.627673

**Authors:** Mitsuhiro Kimura, Takeshi Yoshizumi

**Affiliations:** Faculty of Agriculture, Takasaki University of Health and Welfare, Takasaki-shi, Gunma 370- 0033, Japan

**Keywords:** leaf lettuce, shoot regeneration efficiency, 6-benzylaminopurine (BAP), seed coat color, CIELAB color scale

## Abstract

Lettuce (*Lactuca sativa* L.) is an economically important leafy vegetable crop around the world. Recently, transgenic lettuce has been developed to improve agronomic traits. Although plant regeneration is a key step in plant genetic transformation, it depends strongly on the cultivar. In this study, shoot regeneration efficiency was measured by using two leaf lettuce cultivars, ‘Chima-sanchi’, white seed cultivar, and ‘Chirimen-chisya’, brown seed cultivar, a basal salt mixture, and a combination of the cytokinin, 6-benzylaminopurine (BAP), and an auxin, 1- naphthaleneacetic acid (NAA). Efficient shoot regeneration in ‘Chima-sanchi’ was thus obtained on 1 x Murashige and Skoog medium (MS) with 0.05 mg L^-1^ BAP and 0.1 mg L^-1^ NAA. However, the highest efficiency was obtained in ‘Chirimen-chisya’ on 1 x MS with 0.5 mg L^-1^ BAP and 0.1 mg L^-1^ NAA. Therefore, the BAP concentration for efficient shoot regeneration differs significantly between the two cultivars. These cultivars also have different seed coat colors. Due to the similar results obtained from analyzing 4 additional cultivars, 2 white seed cultivars and 2 brown seed cultivars, we demonstrated a strong correlation between the BAP concentration for efficient shoot regeneration and the seed coat color in leaf lettuce.

## Introduction

Lettuce (*Lactuca sativa* L.), originating in the Mediterranean area, is a major commercial salad crop of the *Asteraceae* family. According to the Food and Agriculture Organization of the United Nations (FAO), total world production of lettuce and chicory has increased by 1.5-fold over the last 20 years [1]. In 2020, lettuce production in Asia was 17.4 million tons, or 63% of global production; in Japan production was 0.6 million tons, sixth highest in the world [1]. Lettuce is remarkably useful as a dietary source of vitamins and minerals [2]. Thus, lettuce cultivars with improved yield and resistance to biotic and abiotic stresses have been developed by conventional breeding methods [3,4].

Recently, transgenic and transplastomic lettuce lines that not only improve agronomic traits but also accumulate pharmaceutical proteins and bio-compounds have been developed using transformation procedures mediated by particle bombardment and *Agrobacterium* [5–8]. There are 5 major types of lettuce varieties in the world: leaf, crisphead, butterhead, romaine, and stem lettuce [9]. Leaf lettuce varieties feature no head and have wrinkled leaves with frilly edges; their shoot fresh weights are higher than those of the butterhead varieties under light-emitting diode (LED) lighting [10]. Therefore, leaf lettuce varieties are better suited to indoor growing. Moreover, leaf lettuce contains more beta-carotene, a precursor to vitamin A, and more lutein per dry weight than either crisphead or butterhead lettuce [11].

Plant tissue cultures have also been utilized widely in plant breeding and industrial applications, such as for the propagation of virus-free plants, the production of valuable compounds, or somaclonal variations [12]. The shoot regeneration efficiency of most plant species is highly dependent on the explant sources, the basal salt mixtures, sugars, and plant growth regulators (PGRs) [13–22]. Previous reports indicated that a combination of the cytokinin, 6-benzylaminopurine (BAP), and an auxin, 1-naphthaleneacetic acid (NAA) is effective for shoot regeneration of lettuce, but, optimal combination differs among the cultivars [23–26]. The optimization of plant tissue culture parameters is labor-intensive and time-consuming as plant tissue cultures often requires long-term. Recently, the molecular mechanisms regulating shoot regeneration in lettuce have been examined [27]. However, molecular approach to improve shoot regeneration efficiency for each strain and cultivar is very expensive. It has been reported that normal rhizomes in *Cymbidium kanran* Makino have the potential for shoot regeneration, but fascinated rhizomes do not [28]. Although the traits are useful as morphological marker for efficient shoot regeneration, no marker has been identified in lettuce.

In this study, we showed that a strong correlation between BAP concentration for efficient shoot regeneration and seed coat color in leaf lettuce. The marker associated with tissue culture was first identified and may be very useful for breeding and genetic engineering of lettuce.

## Materials and Methods

### Plant materials and growth conditions

Six leaf lettuce cultivars were used for this research: ‘Chima-sanchi’ and ‘Chirimen- chisya’ (purchased from Tohoku Seed, Tochigi, Japan), ‘Red fire’ and ‘Green wave’ (purchased from Takii Seed, Kyoto, Japan), and ‘Fringe green’ and ‘Shiki-beni’ (purchased from Sakata Seed, Kanagawa, Japan). ‘Chima-sanchi’, ‘Red fire’ and ‘Shiki-beni’ are white seed cultivars, and ‘Chirimen-chisya’, ‘Green wave’ and ‘Fringe green’ are brown seed cultivars (Figure 3A). The seeds were stored in the constant humidity chamber (SD-302-01, Toyo Living, Kanagawa, Japan) at room temperature and relative humidity of 0 - 1% until sowing. The seeds were treated with 70% EtOH for 1 min, followed by 20% commercial chlorine bleach (Kao, Tokyo, Japan) for 15 min, then rinsed 3 times with sterile distilled water. The sterilized seeds were placed on a germination medium containing half-strength 1 x Murashige and Skoog medium (MS) medium [29] with 10 g L^-1^ sucrose and 2.5 g L^-1^ Phytagel (Sigma-Aldrich, St. Louis, MO) in 9 cm diameter petri dishes. The pH of the medium was adjusted to 5.8 with 1N KOH, before autoclaving at 120°C, pressure at 0.1 MPa, and duration for 20 minutes. Seeds were germinated in an environmentally controlled growth chamber (LPH-411S, NK systems, Tokyo, Japan) fitted fluorescent light (FLR40SW/M/36, NEC, Tokyo, Japan) at a PPFD of 300 μmol photons m^−2^ s^−1^ under continuous white light (cWL) conditions at 20 °C.

### Media composition on shoot regeneration

Shoot regeneration efficiency was examined by the medium supplemented with different basal salt mixtures, sugars, and concentrations of BAP and NAA (Nacalai Tesque, Kyoto, Japan) based on the method described previously [5] (Table 1). All media contained 0.5 mg L^-1^ polyvinylpyrrolidone (PVP, Nacalai Tesque, Kyoto, Japan), and 2.5 g L^-1^ Phytagel (Sigma-Aldrich, St. Louis, MO, USA) at pH 5.8. The media were sterilized by autoclaving at 121 °C for 20 minutes. The cotyledons from 7-day-old seedlings as explant were placed on the medium in 9cm diameter petri dishes (16 explants per dish). The explants were maintained for 4 weeks under cWL conditions at 25 °C and transferred to fresh medium every 2 weeks.

**Table 1.**
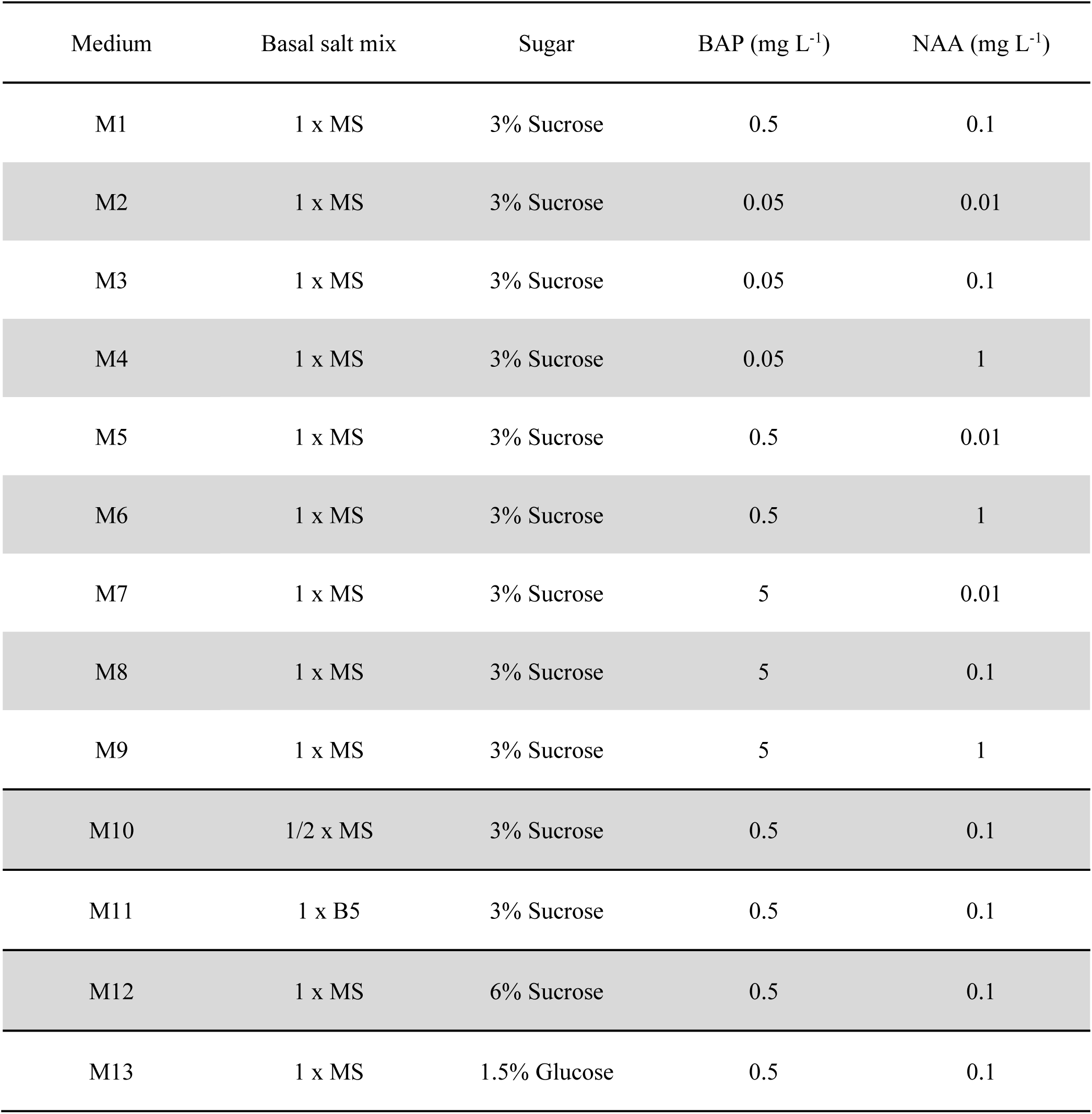
Media composition and growth regulators for shoot regeneration of lettuce.

### Seed coat color measurement

The color parameters of seeds in each cultivar were measured with an SD 7000 spectrophotometer (Nippon Denshoku Industries, Tokyo) using the CIELAB *L\**, *a*,* and *b\** color scale. The *L\** axis represents the degree of brightness from black (*L\** = 0) to white (*L\** = 100). The *a\** and *b\** axes represent redness (positive number) / greenness (negative number) and yellowness (positive number) / blueness (negative number), respectively [30].

### Total flavonoid content analysis

Total flavonoid content was analyzed according to a previous method with some modifications [31]. A 50 µg aliquot of seeds was homogenized in 0.5 mL of 80% methanol with 5.0 mm stainless beads (Biomedical Science, Tokyo, Japan) at 1,100 rpm for 45 s using Shake Master (Biomedical Science, Tokyo, Japan). The homogenized solutions were incubated for 15 min at 70°C and centrifuged at 10,000 xg for 10 min at 4°C. Supernatants were incubated at 60°C, and the dried pellets were dissolved in 20 µL of 80% methanol. The extracts were spotted on a 5 cm x 5 cm TLC Silica gel 60 F₂₅₄ plate (Merck, Darmstadt, Germany). The blots were stained by spraying with a methanolic solution including 1% diphenylboric acid 2- aminoethylester (DPBA, Tokyo Chemical Industry, Tokyo, Japan), followed by spraying with a methanolic solution including 5% PEG 4000 (Nacalai Tesque, Kyoto, Japan). The fluorescence was visualized by an iBright CL1000 imaging system (Thermo Fisher Scientific, Waltham, MA, USA).

### Statistical analyses

All statistical analyses were performed by EZR software [32]. Statistical significance was determined by Student’s *t*-test for two-group comparisons or by one-way ANOVA followed by Tukey’s test for multiple group comparisons. *P*-values < 0.05 were considered statistically significant. All values are expressed as means ± standard errors (SE).

## Results and Discussion

### Effects of medium composition on shoot regeneration

BAP and NAA are commonly used as PGRs in lettuce shoot regeneration [5,23,25,33]. The different concentrations of BAP and NAA for efficient shoot regeneration were examined in ‘Chima-sanchi’ and ‘Chirimen-chisya’ (Figure 1). When different concentrations of BAP were used together with 0.1 mg L^-1^ NAA (M1, M3, and M8), the highest efficiency in ‘Chima-sanchi’ (Figure 1C) was achieved with the medium containing 0.05 mg L^-1^ BAP (M3), whereas in ‘Chirimen-chisya’ (Figure 1D), the highest efficiency was achieved with 0.5 mg L^-1^ BAP (M1). When different concentrations of NAA were tested together with 0.5 mg L^-1^ BAP (M1, M5, and M6), the overall highest efficiency was achieved in ‘Chima-sanchi’ using a medium containing 0.1 mg L^-1^ NAA (M1) (Figure 1C), but in ‘Chirimen-chisya’ the efficiencies using 0.1 mg L^-1^ (M1) and 1 mg L^-1^ NAA (M6) were not significantly different (Figure 1D). In both cultivars, shoots were poorly regenerative in the presence of 5 mg L^-1^ BAP (M7-9; Figure 1). Thus, our results indicated that the BAP significantly influenced shoot regeneration efficiency in leaf lettuce cultivars and that the efficiency depended on the cultivar. The effects of a basal salt mixture and sugar were no difference between both cultivars (Figure S1 and S2).

**Figure 1.**
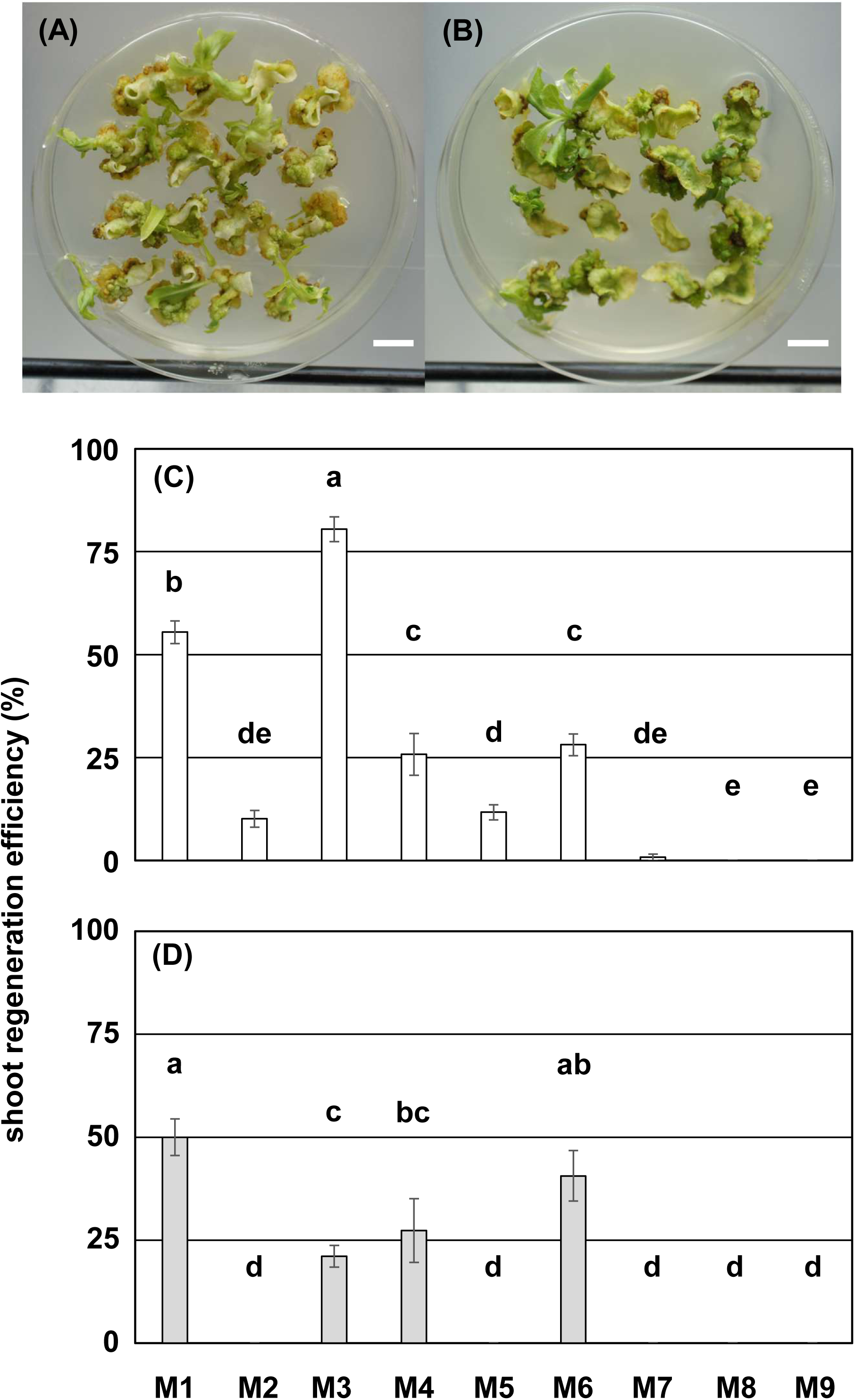
Shoot regeneration from cotyledon segments of ‘Chima-sanchi’ (A) and ‘Chirimen- chisya’ (B) on medium M1 after 4 weeks of culture. Bar = 1 cm. Effects of different concentrations of BAP and NAA on shoot regeneration from cotyledon segments of ‘Chima-sanchi’ (C) and ‘Chirimen-chisya’ (D) after 4 weeks of culture (n = 16 x 8). All media were supplemented with 1 x MS, 30 g L^-1^ Suc, and 500 mg L^-1^ of PVP. Different letters indicate statistically significant differences (one-way ANOVA followed by Tukey’s test, P < 0.05).

### Seed coat-color phenotype of leaf lettuce

Many plant families, e.g., Fabaceae and Poaceae, exhibit various seed coat-color phenotypes, and these phenotypes are important agronomic traits [34–37]. Seed color also has been investigated in lettuce cultivars [38]. In this study, the seed coat-color phenotypes of ‘Chima-sanchi’ and ‘Chirimen-chisya’ were evaluated by measuring seed coat color with the CIELAB color coordinates (*L\**, *a\**, and *b\**) as shown in Figure 2. All CIELAB values were positive for both cultivars, and all CIELAB values of ‘Chima-sanchi’ were higher than those of ‘Chirimen-chisya’, indicating that ‘Chima-sanchi’ seeds were lighter, redder, and yellower than ‘Chirimen-chisya’ seeds (Figure 2B-D).

**Figure 2.**
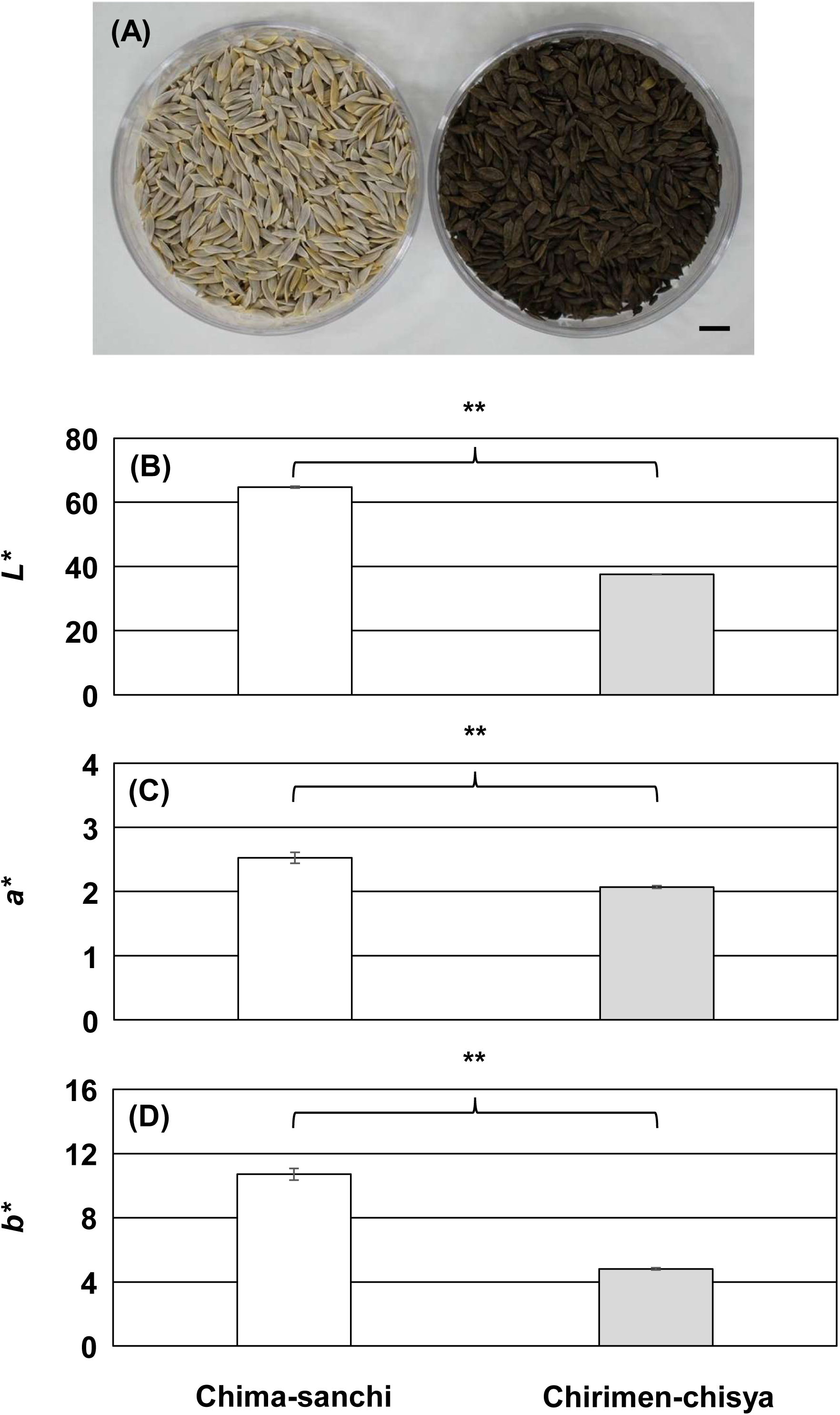
(A) Seed samples of ‘Chima-sanchi’ (left) and ‘Chirimen-chisya’ (right). Average values of CIE *L\** (B), *a\** (C), *b\** (D) color coordinates of seed coat color in the cultivars (n = 5). Horizontal bars within the box indicate the median value of the data, and the outer vertical bars represent the maximum and minimum values of the data. **: P < 0.01 (Student’s *t*-test).

### Relationship between seed coat color and the BAP concentration for efficient shoot regeneration in leaf lettuce

Our results revealed differences in both seed coat color and 0.05 and 0.5 mg L^-1^ BAP for shoot regeneration between ‘Chima-sanchi’ and ‘Chirimen-chisya’ (Figures 1 and 2). Therefore, the relationship between seed coat color and 0.05 and 0.5 mg L^-1^ BAP was further analyzed in 4 additional cultivars: ‘Red fire’, ‘Shiki-beni’, ‘Green wave’, and ‘Fringe green’ (Figure 3A). 0.05 and 0.5 mg L^-1^ BAP was calculated as the ratio of shoot regeneration efficiency using M3 to that using M1 (M3/M1, Table S1). A significantly positive correlation was found between the M3/M1 and the seed coat-color values of *L\** and *b\** (r=0.834 and 0.722) (Figure 3B, D), but no correlation with the *a\** value was found (r=0.004) (Figure 3C). The relationships between seed coat color and other agronomic traits, such as preharvest spouting tolerance and seed impermeability, are well known in several plant species [39,40]. In addition, Arabidopsis *transparent testa* (*tt*) and *transparent testa glabra* (*ttg*) mutants have been isolated by examining coat colors and permeability [41]. Nameth et al. (2013) reported that the Arabidopsis *tt4* mutant is deficient in chalcone synthase (CHS), which is involved in the flavonoid biosynthesis pathway, shows a pale yellow seed color, and has significantly low shoot regeneration efficiency. Thus, some downstream flavonoid after catalysis by CHS might be required for efficient shoot regeneration in Arabidopsis. In this study, brown seed cultivars in leaf lettuce contained more flavonoid than white seed cultivars (Figure S3). However, the shoot regeneration efficiencies of white seed cultivars were much higher than those of brown seed cultivars (Figure 1). Although shoot regeneration in leaf lettuce and Arabidopsis might be regulated by several different mechanisms, seed coat color has a strong correlation to optimize BAP concentration for efficient shoot regeneration in leaf lettuce.

**Figure 3.**
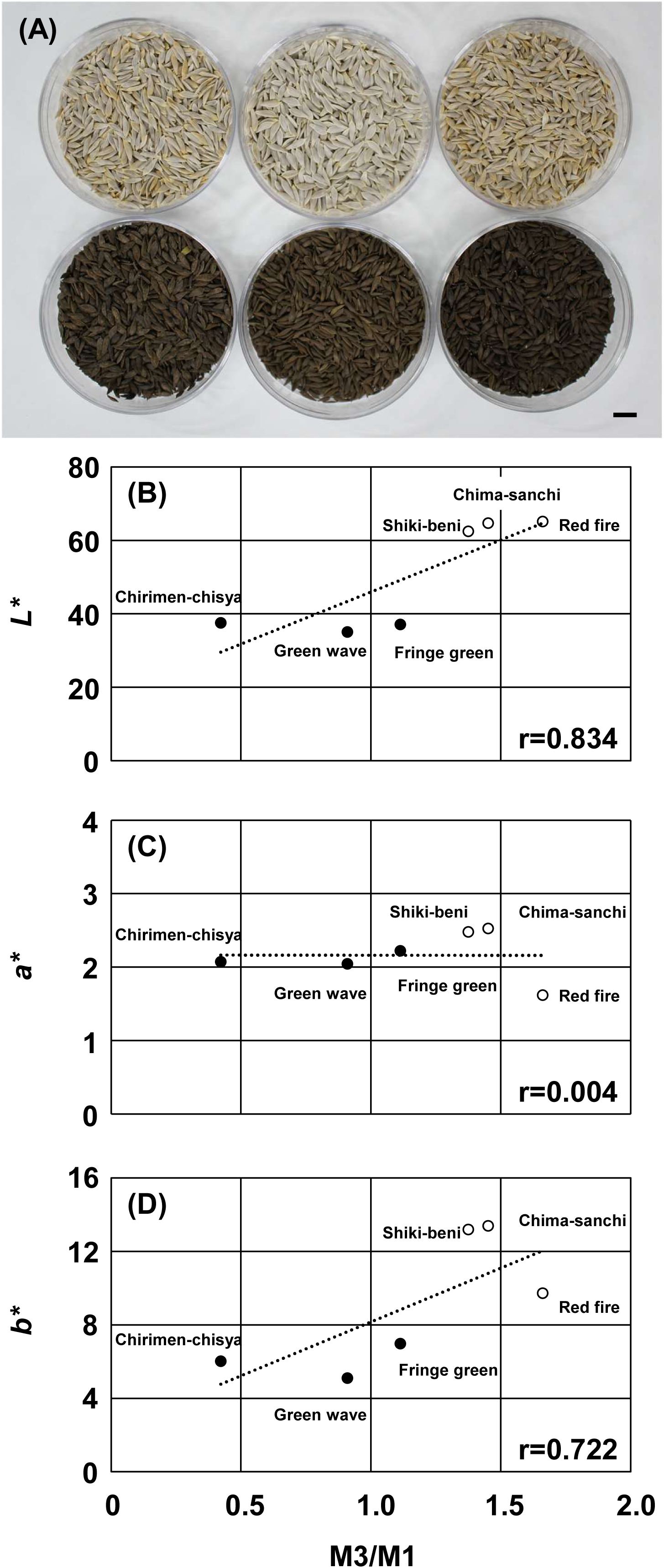
(A) Samples of six lettuce cultivars. Genotype names are as follows, clockwise from the upper left: chima-sanchi, red fire, shiki-beni, green wave, fringe green, and chirimen-chisya. Bar = 1.0 cm. Correlations between values of CIE *L\** (B), *a\** (C), *b\** (D) color coordinates of seed coat color (n = 5) and ratio of shoot regeneration efficiency using M7 to the efficiency using M1 (M7/M1) in the 6 cultivars (n = 16 x 8).

## Conclusion

This study revealed the relationship between BAP concentration for efficient shoot regeneration and seed coat color in leaf lettuce. The BAP concentration for efficient shoot regeneration in white seed cultivars was 0.05 g L^-1^, whereas it was 0.5 g L^-1^ in brown seed cultivars. There was not significantly difference in basal salt mixture, sugar, or concentration of NAA between white and brown seed cultivars. Our results will support the further study of shoot regeneration in other lettuce cultivars and related species.

### Author contribution

M.K. and T.Y. conceived, designed, and supervised the project. M.K. performed all experiments. T.Y. provided some important suggestions. M.K. wrote the paper. T.Y. reviewed and commented on the manuscript.

## Funding

This work was supported by JSPS KAKENHI Grant Number JP18K05638 (M. K.) and 20K21319 (T. Y), New Energy and Industrial Technology Development Organization (NEDO) Grant (T. Y.) and the Research Grant in Takasaki University of Health and Welfare (M. K. and T. Y.).

### Data Availability Statement

All data are available with an email request to the authors.

## Acknowledgements

We are grateful to Ms. Ayumi Nagai (Takasaki University of Health and Welfare), Mr. Ryoei Kawakami, and Ms. Ayumi Yamato (Gunma Industrial Technology Center) for their technical support, Dr. Akira Endo (Kaneka Corporation) and Prof. Eiji Nambara (University of Toronto) for their critical suggestions.

