## supplemental figure and table for "Relationship between concentration of cytokinin for efficient shoot regeneration and seed coat color in leaf lettuce"

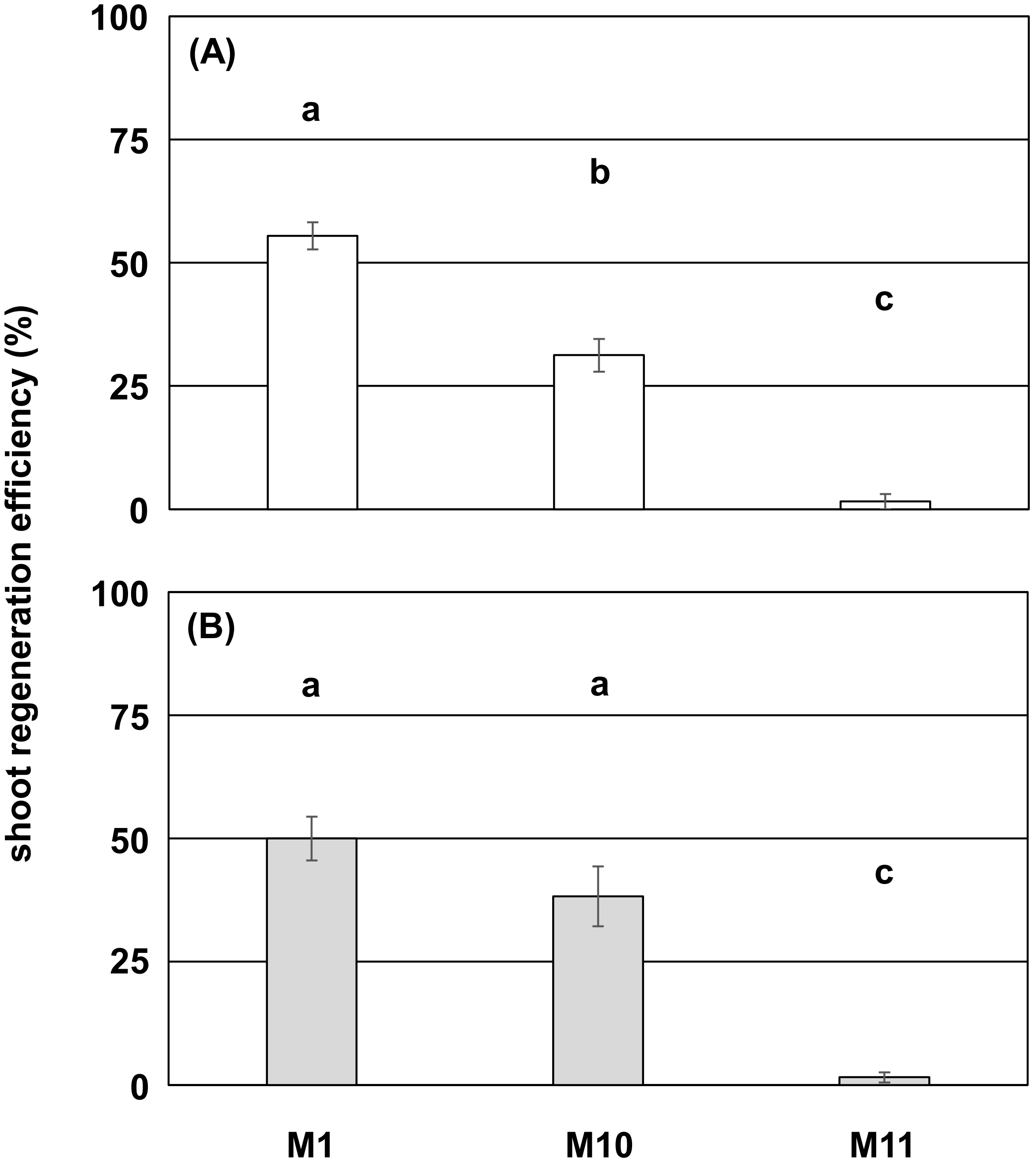


**Figure S1** Effects of different basal media on shoot regeneration from cotyledon segments of 'Chima-sanchi' (A) and 'Chirimen-chisya' (B) after 4 weeks of culture (n = 16 x 8). All media were supplemented with 30 g $L^{-1}$ Suc, 0.5 mg $L^{-1}$ BAP, 0.1 mg $L^{-1}$ NAA, and 500 mg $L^{-1}$ PVP. Different letters indicate statistically significant differences (one-way ANOVA followed by Tukey's test, $P < 0.05$).

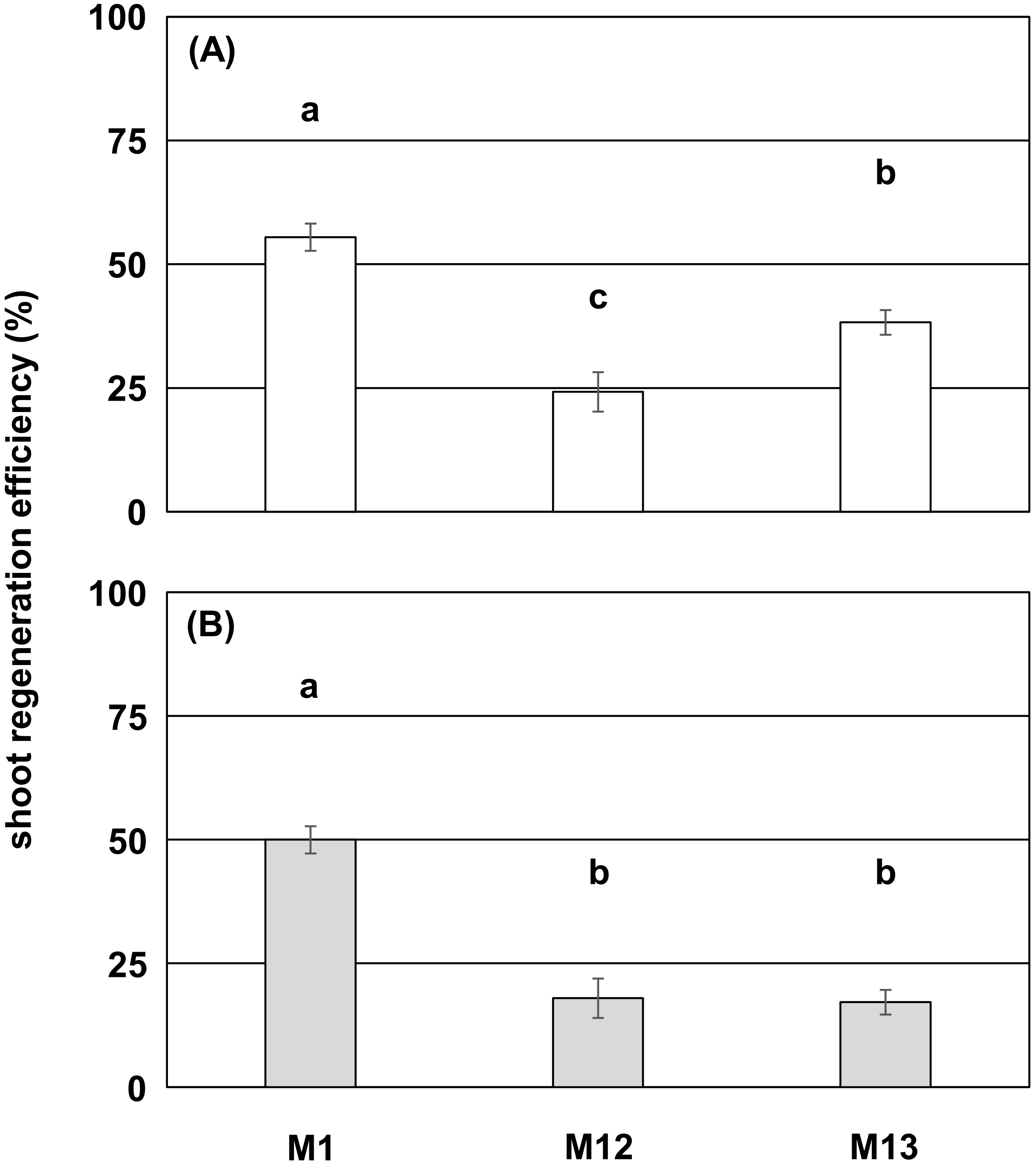


**Figure S2** Effects of different sugar on shoot regeneration from cotyledon segments of 'Chima-sanchi' (A) and 'Chirimen-chisya' (B) after 4 weeks of culture (n = 16 x 8). All media were supplemented with 1 x MS, 0.5 mg $L^{-1}$ BAP, 0.1 mg $L^{-1}$ NAA, and 500 mg $L^{-1}$ of PVP. Different letters indicate statistically significant differences (one-way ANOVA followed by Tukey's test, $P < 0.05$).

Chima-sanchi Red fire Shiki-beni

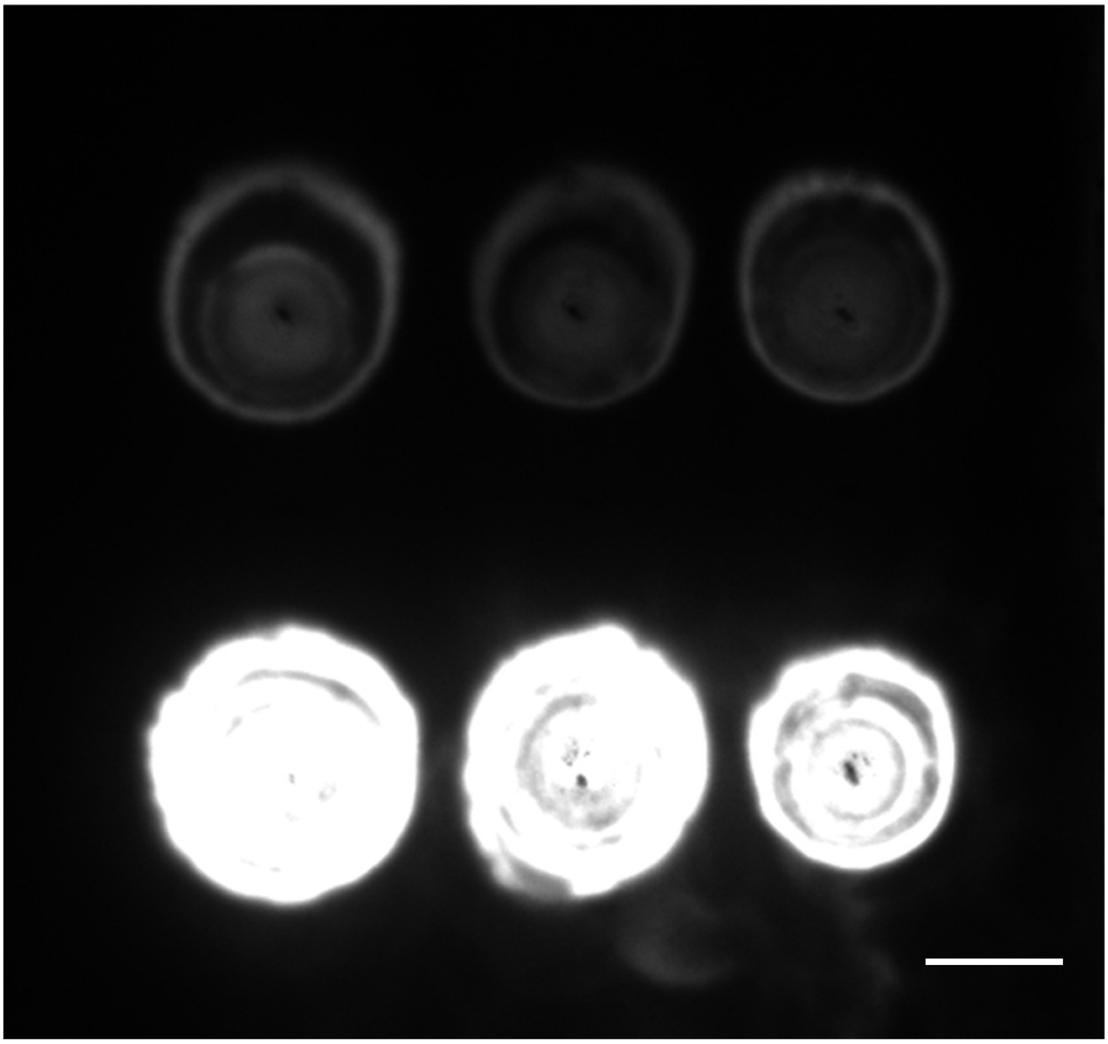

Chirimen-chisya Fringe green Green wave

**Figure S3** The flavonoid content of methanolic extracts of seeds in 6 lettuce cultivars. Bar = 0.5 cm

**Table S1** Shoot regeneration efficiency using M1 and M3 and CIE Lab values of seed coat color in 6 lettuce cultivars

| **Cultivar[1]** | **Shoot regeneration efficiency (%)** | | | | | **CIE Lab values[2]** | | | | | |
|---|---|---|---|---|---|---|---|---|---|---|---|
| | **M1** | | **M3** | | **M3/M1** | ***L**** | | ***a**** | | ***b**** | |
| | **Mean** | **SE** | **Mean** | **SE** | | **Mean** | **SE** | **Mean** | **SE** | **Mean** | **SE** |
| **Chima-sanchi (w)** | **55.47** | **2.75** | **80.47** | **3.00** | **1.45** | **64.72** | **0.31** | **2.53** | **0.09** | **13.39** | **0.45** |
| **Red fire (w)** | **46.09** | **2.62** | **76.56** | **3.69** | **1.66** | **65.19** | **0.07** | **1.62** | **0.02** | **9.73** | **0.06** |
| **Shiki-beni (w)** | **56.25** | **5.28** | **77.34** | **2.88** | **1.36** | **62.52** | **0.05** | **2.48** | **0.05** | **13.19** | **0.16** |
| **Chirimen-chisya (b)** | **50.00** | **4.42** | **21.09** | **2.62** | **0.42** | **37.57** | **0.09** | **2.07** | **0.02** | **6.01** | **0.08** |
| **Fringe green (b)** | **41.41** | **4.72** | **46.09** | **3.53** | **1.11** | **37.15** | **0.03** | **2.22** | **0.02** | **6.97** | **0.04** |
| **Green wave (b)** | **68.75** | **4.26** | **62.50** | **2.64** | **0.91** | **35.08** | **0.09** | **2.05** | **0.01** | **5.10** | **0.03** |

**[1]: w; white seed cultivar, b; brown seed cultivar**

**[2]: *L** corresponding to the brightness, *a** to the red/green coordinates and *b** to the yellow/blue coordinates.**
